# Mediator subunit MDT-15 promotes expression of propionic acid breakdown genes to prevent embryonic lethality in *Caenorhabditis elegans*

**DOI:** 10.1101/2022.12.16.520805

**Authors:** Grace Ying Shyen Goh, Arshia Beigi, Junran Yan, Kelsie R. S. Doering, Stefan Taubert

**Author notes:** **GYSG**: Semios Biotechnologies, 3430 Brighton Ave #204A, Burnaby, BC V5A 3H4; **AB**: Department of Medicine, University of British Columbia, 2775 Laurel Street Vancouver, British Columbia Canada V5Z 1M9. **Corresponding author:** Stefan Taubert, Centre for Molecular Medicine and Therapeutics, Room 2024, 950 W 28th Avenue, Vancouver, BC, Canada, V5Z 4H4.

## Abstract

The micronutrient vitamin B12 is an essential cofactor for two enzymes: methionine synthase, which plays a key role in the one-carbon cycle; and methylmalonyl-CoA mutase, an enzyme in a pathway that breaks down branched-chain amino acids and odd-chain fatty acids. A second, vitamin B12-independent pathway that degrades methylmalonyl-CoA and its upstream metabolite propionic acid was recently described in *Caenorhabditis elegans*, the propionate shunt pathway. Activation of five shunt pathway genes in response to low vitamin B12 availability or high propionic acid levels is accomplished by a transcriptional regulatory mechanism involving two nuclear hormone receptors, NHR-10 and NHR-68. Here, we report that the *C. elegans* Mediator subunit *mdt-15* is also essential for the activation of the propionate shunt pathway genes, likely by acting as a transcriptional coregulator for NHR-10. *C. elegans mdt-15* mutants fed a low vitamin B12 diet have transcriptomes resembling those of wild-type worms fed a high vitamin B12 diet, with low expression of the shunt genes. Phenotypically, the embryonic lethality of *mdt-15* mutants is specifically rescued by diets high in vitamin B12, but not by dietary polyunsaturated fatty acids, which rescue many other phenotypes of the *mdt-15* mutants. Finally, NHR-10 binds to MDT-15 in yeast-two-hybrid assays, and the transcriptomes of *nhr-10* mutants resemble those of *mdt-15* mutants. Our data show that MDT-15 is a key coregulator for an NHR regulating propionic acid detoxification, adding to roles played by NHR:MDT-15 partnerships in metabolic regulation and pinpointing vitamin B12 availability as a requirement for *mdt-15* dependent embryonic development.

## Introduction

Animals adjust their metabolism based on their nutritional environment, which allows them to optimize the use of available resources and adjust to or compensate for absent or limiting nutrients. Such metabolic adjustments occur through various mechanisms, including alteration in the expression of metabolic and other genes via transcriptional modulation. Regulation can involve direct sensing of nutritional components or metabolites by transcription factors such as nuclear hormone receptors (NHRs) and indirect sensing via upstream regulators such as G protein-coupled receptors or other sensors (Schneider-Poetsch and Yoshida 2014; Boukouris *et al.* 2016; Husted *et al.* 2017; Suganuma and Workman 2018).

Vitamins are a group of essential micronutrients required in many developmental and physiological functions. All animals require vitamin B12, which is vital for DNA synthesis, and fatty acid and amino acid metabolism (Deodato *et al.* 2017; Froese *et al.* 2019). Vitamin B12 is an essential cofactor for methylmalonyl-CoA mutase (MUT), an enzyme in the canonical and evolutionarily conserved propionic and methylmalonic acid breakdown pathway (Bito and Watanabe 2016; Deodato *et al.* 2017; Froese *et al.* 2019). Loss of activity in this pathway due to mutations in the MUT gene or in the genes for propionyl-CoA-carboxylase (PCCA/B) causes the childhood diseases methylmalonic acidemia and propionic acidemia, respectively (Deodato *et al.* 2017). Abnormal metabolism in these genetic diseases severely affects development and neurological functions and can cause lethality in newborns.

Recent work in *Caenorhabditis elegans* has revealed the existence of a parallel, vitamin B12-independent propionic and methylmalonic acid breakdown pathway, termed the propionic and methylmalonic acid breakdown shunt (‘the shunt’) (Watson *et al.* 2016). The shunt is composed of enzymes that are conserved in humans (*acdh-1/ACABCB, ech-6/ECHS1, hach-1/HIBCH, hpdh-1/ADHFE1, alh-8/ALDH6A1*) (Watson *et al.* 2016). However, as intermediate metabolites of the shunt are toxic, shunt gene expression is repressed except when the pathway is absolutely required. In *C. elegans*, conditions of low dietary vitamin B12 or high dietary propionic acid result in increased shunt gene expression (MacNeil *et al.* 2013; Watson *et al.* 2013, 2014, 2016).

The tight regulation of shunt genes in *C. elegans* is achieved by combinatorial activity of NHR-10 and NHR-68 (Bulcha *et al.* 2019). Both NHRs are required to induce shunt gene expression and for organismal survival on diets with high propionic acid levels. Interestingly, NHR-10 itself activates *nhr-68*, yet *nhr-68* expression alone is insufficient to induce the propionate shunt genes; rather, *nhr-10* and *nhr-68* are both required for this activation, and hence form a self-reinforcing regulatory circuit that only results in gene activation when elevated propionic acid levels persist in the media for several hours (Bulcha *et al.* 2019).

Like all transcription factors, NHRs do not act in isolation but require coregulators to control gene expression. In *C. elegans*, the Mediator subunit MDT-15 is a coregulator for many NHRs that regulate nutritional and stress adaptive responses (Grants *et al.* 2015; Hartman *et al.* 2021). Accordingly, *mdt-15* mutation or depletion results in many developmental and physiological phenotypes, many of which can be rescued by supplementation with unsaturated fatty acids (Taubert *et al.* 2006; Yang *et al.* 2006; Hou *et al.* 2014; Lee *et al.* 2015, 2019). However, some phenotypes of the *mdt-15* mutant cannot be rescued by unsaturated fatty acids, suggesting that other dietary processes and metabolic regulatory pathways may be linked to *mdt-15* (Goh *et al.* 2014, 2018).

Here, we show that transcriptomic changes caused by *mdt-15* loss resemble those of worms grown on a low vitamin B12 diet, with shunt genes downregulated in both conditions. Supplementation with vitamin B12 rich diets, but not with unsaturated fatty acids, rescued the embryonic lethality of *mdt-15* mutants. NHR-10, which controls shunt gene expression, physically interacts with MDT-15, and loss of either regulator produces similar transcriptomic perturbations. Our data suggest a model wherein *C. elegans* MDT-15 acts as a coregulator for NHR-10, with the two factors inducing shunt genes on demand to ensure propionic acid detoxification and prevent adverse organismal phenotypes.

## Materials and methods

### *C. elegans* growth conditions

We cultured *C. elegans* strains using standard techniques on nematode growth media (NGM) plates, as described (Brenner 1974). NGM plates were supplemented at the indicated concentrations with methylcobalamin (Sigma M9756), adenosylcobalamin (Sigma C0884), propionic acid (Sigma 402907), tBOOH (Sigma 458139), and PUFAs (mix of fatty acid sodium salts: 150 μM C18:2, S-1127; 150 μM C20:5, S-1144, Nu-Chek Prep). *E. coli* OP50 and *C. aquatica* DA1877 were used as food sources, as indicated. *fat-6* RNAi experiments were performed with *E. coli* HT115 on nematode growth media (NGM) plates supplemented with 100 μg/ml carbenicillin (BioBasic CDJ469), 1 mm IPTG (Santa Cruz, sc-202185B), and 12.5 μg/ml tetracycline (BioBasic TB0504). RNAi clones were from the Ahringer (Source BioScience) and were sequenced prior to use. All experiments were carried out at 20°C.

### *C. elegans* strains

Worm strains used in this study were N2 WT and XA7702 *mdt-15(tm2182)* (Taubert *et al.* 2008); the mutant was backcrossed into our lab N2 strain prior to study. For synchronized worm growths, we isolated embryos by standard sodium hypochlorite treatment. Isolated embryos were allowed to hatch overnight on unseeded NGM plates until the population reached a synchronized state of halted development at L1 stage via short-term fasting (16–24 hr). Synchronized L1 stage larvae were then transferred to seeded plates and grown to the desired stage.

### Embryo viability assays

WT N2 and XA7702 *mdt-15(tm2182)* worms were grown to young adults on NGM plates seeded with *E. coli* OP50 or *C. aquatica* DA1877. Then, 10 worms were transferred onto plates seeded with *E. coli* OP50 or *C. aquatica* DA1877 and left at 20°C for 24 hours. Adults were then removed, and progeny were allowed to hatch for 48 hours. Eggs that had not hatched thereafter were classified as non-viable.

### Oxidative stress sensitivity assays

To assess oxidative stress sensitivity, synchronized N2 and *mdt-15(tm2182)* L1 stage worms were allowed to grow on NGM plates containing 300 μM PUFAs and/or 5 nM vitamin B12 until they reached the mid-L4 stage. Then, they were transferred to plates also containing 2 or 4 mM tBOOH, which were seeded with heat-inactivated *E. coli* OP50. After 24 hours, the number of dead or alive worms was counted. In parallel, to ascertain PUFA effectiveness, we also scored whether PUFA supplementation was able to rescue viability of N2 on RNAi plates seeded with HT115 bacteria containing control empty vector (EV) of *fat-6* RNAi clones.

### RNA isolation and RT-qPCR analysis

Synchronized L1 worms were allowed to grow on *E. coli* OP50 plates for 48 hr to L4 stage and rapidly harvested. RNA isolation was performed as previously described (Doering *et al.* 2022). Briefly, 2 μg total RNA was used to generate cDNA with Superscript II reverse transcriptase (Invitrogen 18064-014), random primers (Invitrogen 48190-011), dNTPs (Fermentas R0186), and RNAseOUT (Invitrogen 10777-019). Quantitative PCR was performed in 10 μl reactions using Fast SYBR Master Mix (Life Technologies 4385612), 1:10 diluted cDNA, and 5 μM primer, and an analyzed with an Applied Biosystems StepOnePlus machine. We analyzed the data with the ΔΔCt method. For each sample, we calculated normalization factors by averaging the (sample expression)/(average reference expression) ratios of three normalization genes, *act-1, tba-1*, and *ubc-2.* The reference sample was wild type grown on *E. coli* OP50. We used one-way or two-way ANOVA to calculate statistical significance of gene expression changes and corrected for multiple comparisons using the Tukey method. Primers were tested on serial cDNA dilutions and analyzed for PCR efficiency prior to use. All data originate from three or more independent biological repeats, and each PCR reaction was conducted in technical duplicate. Sequences of RT-qPCR primers are:

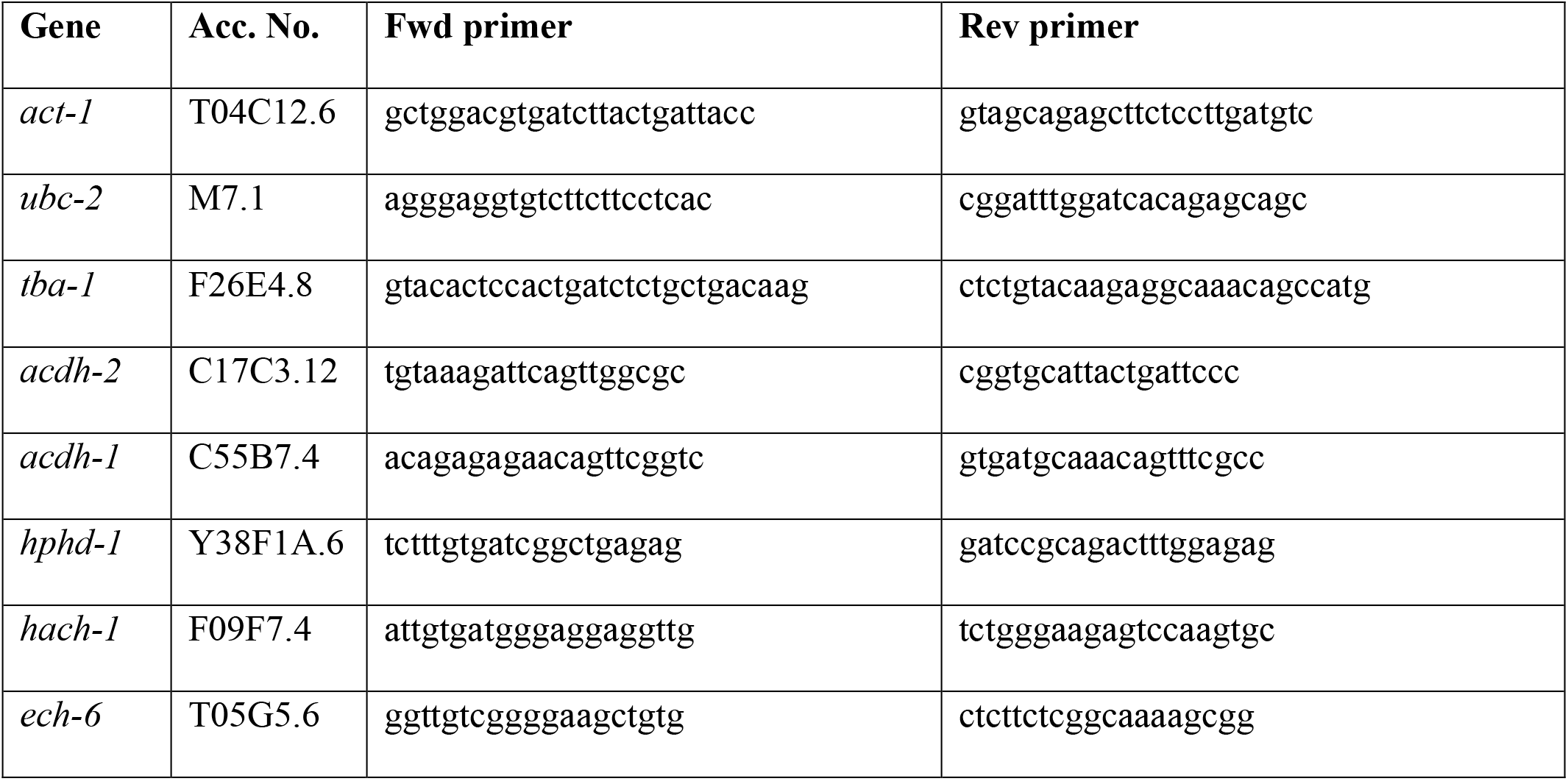

### Microarray analysis

Microarray gene expression profiling of *mdt-15(tm2182)* mutants was performed at the University of California San Francisco SABRE Functional Genomics Facility exactly as previously described (Grants *et al.* 2016), using the same growth conditions, and RNA extraction, processing, hybridization, and analysis methods. Microarray data of *mdt-15(tm2182)* mutants and corresponding WT are deposited in Gene Expression Omnibus (GEO) under a new accession number (GSE220955), with the WT control exactly the same as previously published (GEO Series GSE68520 (Grants *et al.* 2016)). Microarray data were processed using limma (Ritchie *et al.* 2015) exactly as described (Grants *et al.* 2016). To maximize comparability of the microarray to RNA-seq datasets, probe identifiers were converted to gene symbols using the ID conversion module of easyGSEA in the eVITTA toolbox (Cheng *et al.* 2021), and only the highest expressing probe (based on aveA average expression) was kept for each unique gene symbol. Using this approach, we identified 1616 unique differentially expressed genes (DEGs) with Adj.P <0.05 in *mdt-15(tm2182)* mutants, including 563 downregulated (logFC<0) and 1053 upregulated (logFC>0) genes (Supplementary Tables S1-S2).

### Re-analysis of published microarray data

To maximize comparability with our microarray samples, we reanalyzed published two-channel microarray data of *mdt-15(RNAi)* worms (Taubert *et al.* 2008). Raw data was extracted from GEO under the accession number GSE9720 and reanalyzed with limma. Adaptive background correction was performed with the method “normexp” with offset 50, and subsequently normalizeWithinArrays using default method (print tip loess). The command modelMatrix with set reference level “control RNAi” was used to make the sign of M consistent across repeats regardless of dye swapping. Differentially expressed genes were computed using lmFit, eBayes, and topTable with the default parameters, and probe identifiers were converted to unique gene symbols as above. We identified 2451 DEGs with PValue<0.05, including 1298 downregulated and 1153 upregulated genes (Supplementary Tables S1, S3).

We also reanalyzed the published single channel microarray of *Comamonas* DA1877 *vs. E. coli* HT115 treatment (MacNeil *et al.* 2013), for which normalized count data are available in GEO (Accession number: GSE43959). We performed data extraction and DE analysis using easyGEO in the eVITTA toolbox (Cheng *et al.* 2021), using default parameters (limma with quantile normalization and linear model fitting with ls). Probe identifiers were converted to unique gene symbols as above. We identified 6541 DEGs with PValue<0.05, including 2979 downregulated and 3562 upregulated genes (Supplementary Tables S1, S4).

### Re-analysis of published RNA-seq data

To maximize comparability with our microarray samples, we reanalyzed published RNA-sequencing data of Vitamin B12-treated WT worms, and of *nhr-10* and *nhr-68* mutants (Bulcha *et al.* 2019). Relevant SRA accession numbers and metadata were obtained from SRA run selector under the GEO accession number GSE123507. For each SRA accession number, raw reads were downloaded from SRA using prefetch and FASTQ files were extracted using fastq-dump. Because the study was done on BGI-seq-500 platform, we compiled a list of adaptor sequences using the overrepresented sequences from FastQC. Following this, the reads were trimmed using Trimmomatic version 0.36 (Bolger *et al.* 2014) with parameters LEADING:3 TRAILING:3 SLIDINGWINDOW:4:15 MINLEN:36. Next, trimmed reads were aligned to the NCBI reference genome WBcel235 WS277 (https://www.ncbi.nlm.nih.gov/assembly/GCF_000002985.6/) using Salmon version 0.9.1 (Patro *et al.* 2017) with parameters -l A --gcBias --validateMappings. Then, transcript-level read counts were imported into R and summed into gene-level read counts using tximport (Soneson *et al.* 2016). Genes not expressed at a level greater than one count per million (CPM) reads in at least two of the samples were excluded from further analysis. The gene-level read counts were normalized using the trimmed mean of M-values (TMM) in edgeR (Robinson *et al.* 2010) to adjust samples for differences in library size. Differential expression analysis was performed using the quasi-likelihood F-test with the generalized linear model (GLM) approach in edgeR (Robinson *et al.* 2010). The number of upregulated and downregulated DEGs are found in Supplementary Table S1 and all DEGs are listed in Supplementary Tables S5-S7.

### Comparison and visualization of transcriptome data

As quality controls, we compared genes deregulated in *mdt-15(tm2182)* mutants to genes deregulated in *mdt-15(RNAi)* worms (GSE9720), revealing a substantial correlation between the two datasets (Supplementary Fig. S1A), as expected. The scatter plot was generated using plotly and Pearson correlation coefficient was computed using the cor.test function.

To visualize the downregulation of shunt genes in different datasets, we generated volcano plots using limma function plotWithHighlights. We separated differentially expressed genes (DEGs) (defined as the genes with Adj.P or PValue<0.05) above the line, and shunt genes were marked using additional text labels.

To compare our microarray data to previously published data, we generated Venn diagrams using the R package eulerr (Larsson and Gustafsson 2018). Hypergeometric p-value was calculated using the phyper function, where q = size of overlap-1, m = number of genes in gene list 1, n = platform size - m, k = number of genes in gene list 2. For each pairwise comparison, platform size was calculated as the total number of genes that are detected in DE analysis in either sample.

## Results

### Transcriptomes of *mdt-15* mutants resemble those of worms grown on *Comamonas aquatica*

We and others found that *mdt-15(RNAi)* and *mdt-15(tm2182)* hypomorph mutants display phenotypes such as embryonic lethality, delayed larval development, reduced body size, reduced fecundity, fat storage defects, axon migration defects, stress sensitivity, impaired locomotion, and a short lifespan (Taubert *et al.* 2006, 2008; Yang *et al.* 2006; Arda *et al.* 2010; Steimel *et al.* 2013; Goh *et al.* 2014, 2018; Pukkila-Worley *et al.* 2014; Lee *et al.* 2015, 2019; Vozdek *et al.* 2018; Peterson *et al.* 2019; Shomer *et al.* 2019; Doering *et al.* 2022). *mdt-15* inactivation compromises fatty acid desaturation, and PUFA supplementation of worms with reduced *mdt-15* function improves many of the above phenotypes, robustly rescuing larval development, body size, locomotion, fecundity, and life span (Taubert *et al.* 2006; Yang *et al.* 2006; Lee *et al.* 2015, 2019). In contrast, the oxidative stress sensitivity and altered zinc storage of *mdt-15* mutants are not rescued by PUFA supplementation (Goh *et al.* 2014; Shomer *et al.* 2019). Similarly, the embryonic lethality of *mdt-15(tm2182)* hypomorph mutants is unaffected by PUFA supplementation (see below), suggesting that other, unknown dysregulated processes must underlie this phenotype.

To identify *mdt-15* dependent processes that may promote PUFA-independent embryonic development, we compared transcriptome profiles of *mdt-15(RNAi)* and *mdt-15(tm2182)* hypomorph mutants to other transcriptomes. We found that the transcriptomes of wild-type worms fed the bacterial food source *Comamonas aquatica* DA1877 (MacNeil *et al.* 2013) shared a statistically significant overlap with genes dependent on *mdt-15* (Fig. 1A). Vitamin B12 (aka cobalamin) is a key molecule that drives *C. aquatica*-induced developmental acceleration and it is virtually undetectable in *E. coli* OP50 (Watson *et al.* 2014). Accordingly, the transcriptomes of worms grown on vitamin B12 supplemented OP50 (Bulcha *et al.* 2019) also shared a statistically significant overlap with genes dependent on *mdt-15* (Fig. 1B). Thus, loss of *mdt-15* results in transcriptomes that partially resemble those of worms fed vitamin B12 rich diets.

**Figure 1.**
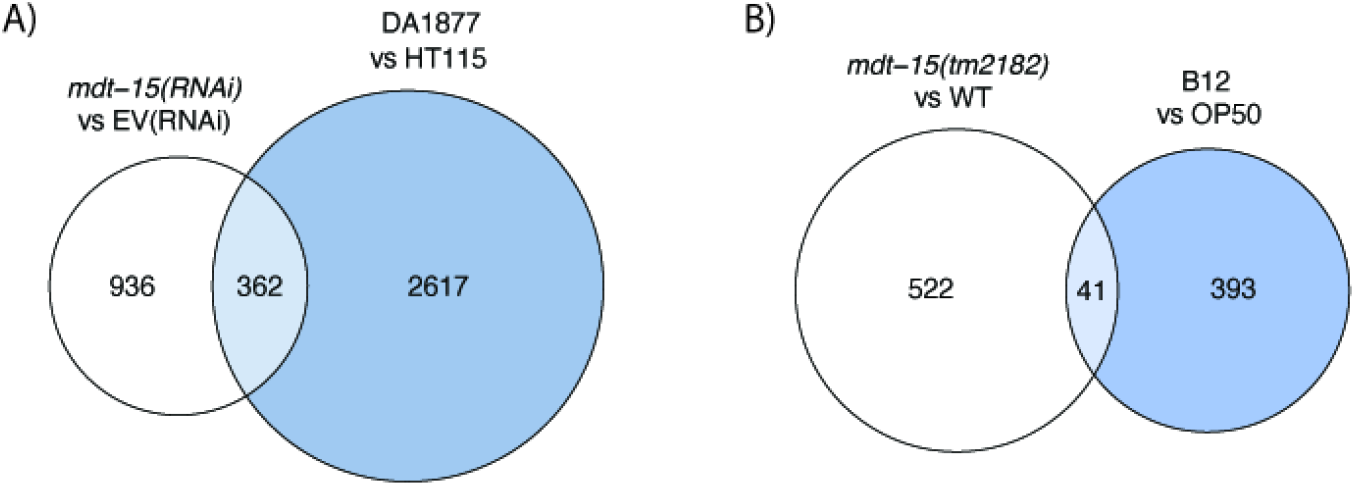
Genes regulated by *mdt-15* overlap with genes regulated by vitamin B12 rich diets. **(A)** The Venn diagram shows overlaps of genes downregulated in *mdt-15(RNAi)* worms and in worms grown on vitamin B12 rich *C. aquatica* DA1877 (PValue<0.05 and logFC<0). Hypergeometric p = 4.37e-28; platform size = 18286. **(B)** The Venn diagram shows overlaps of genes downregulated in *mdt-15(tm2182)* worms and in worms grown on an *E. coli* OP50 diet supplemented with vitamin B12 (Adj.P or PValue<0.05 and logFC<0). Hypergeometric p = 1.31e-10; platform size = 18536. Lists of genes in each data set are shown in Supplementary Tables S2-5.

### A diet of *C. aquatica* rescues the embryonic lethality of *mdt-15(tm2182)* mutants

To determine if a diet of *C. aquatica* DA1877 might influence the phenotypes of *mdt-15(tm2182)* mutants, we compared *mdt-15(tm2182)* mutants fed *C. aquatica* DA1877 or *E. coli* OP50. We observed a substantial increase in the number of viable *mdt-15(tm2182)* mutant offspring when provided *C. aquatica* DA1877 (Fig. 2A). To determine whether maternal or offspring food source caused the phenotypic rescue, we grew *mdt-15(tm2182)* mutants to adulthood on either *E. coli* OP50 or *C. aquatica* DA1877, switched them to the identical or reciprocal food sources, and then quantified the number of surviving offspring. We found that adult *mdt-15(tm2182)* mutants raised on *E. coli* OP50 generated mostly arrested progeny even when switched to *C. aquatica* DA1877, whereas the progeny of worms raised on *C. aquatica* DA1877 mostly hatched successfully even when the adults were switched to an *E. coli* OP50 diet (Fig. 2B). Thus, maternal *C. aquatica* DA1877 is sufficient to rescue the embryonic lethality in *mdt-15(tm2182)* progeny.

**Figure 2.**
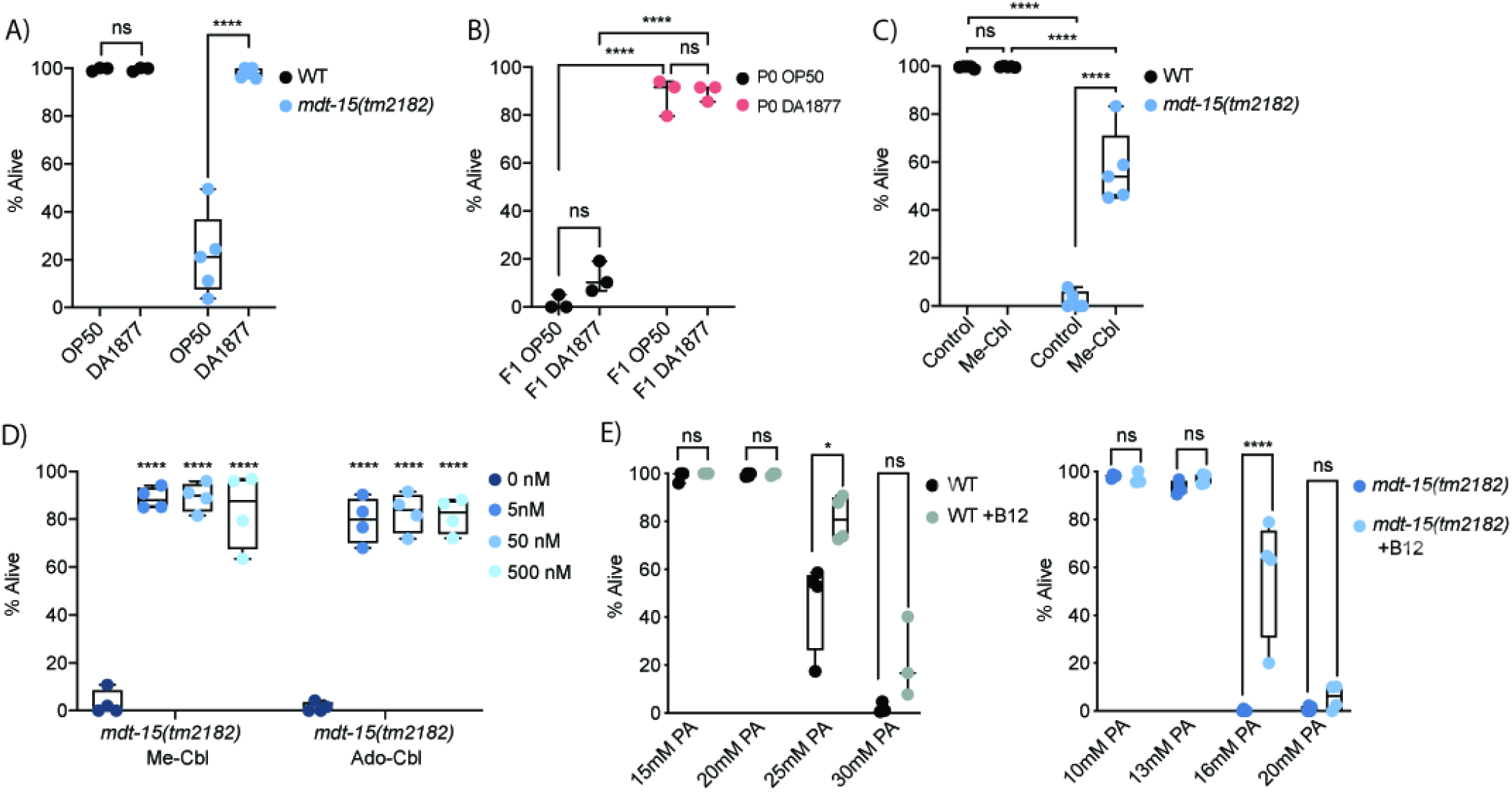
Diets rich in vitamin B12 rescue the embryonic lethality of *mdt-15(tm2182)* mutants. **(A)** The graph shows the average embryonic lethality of wild-type and *mdt-15(tm2182)* mutant worms grown on either *E. coli* OP50 or *C. aquatica* D1877 (n=3-5). ****p<0.001 for indicated comparisons, all other comparisons not significant, as assessed by two-way ANOVA with Tukey correction for multiple comparisons. **(B)** The graph shows the average embryonic lethality of *mdt-15(tm2182)* mutant worms grown on either *E. coli* OP50 or *C. aquatica* D1877 for the P0 and F1 generations, as indicated (n=3). ****p<0.001 for indicated comparisons, all other comparisons not significant, as assessed by two-way ANOVA with Tukey correction for multiple comparisons. **(C)** The graph shows the average embryonic lethality of wild-type and *mdt-15(tm2182)* mutant worms grown on either *E. coli* OP50, either unsupplemented or supplemented with methyl-cobalamin (n=5). ****p<0.001 for indicated comparisons, all other comparisons not significant, as assessed by two-way ANOVA with Tukey correction for multiple comparisons. **(D)** The graph shows the average embryonic lethality of *mdt-15(tm2182)* mutant worms grown on either *E. coli* OP50, either unsupplemented or supplemented with methyl-cobalamin or adenosyl-cobalamin at the indicated concentrations (n=4). ****p<0.001 for indicated comparisons, all other comparisons not significant, as assessed by ordinary one-way ANOVA with Tukey correction for multiple comparisons. **(E)** The graph shows the average embryonic lethality of wild-type (left) or *mdt-15(tm2182)* mutant (right) worms grown on *E. coli* OP50 on the indicated propionic acid concentrations and either unsupplemented or supplemented with methyl-cobalamin, as indicated (n=3-4). *p<0.05, **** p<0.001 for the indicated comparisons, as assessed by two-way ANOVA with Tukey correction for multiple comparisons.

### Vitamin B12 rescues the embryonic lethality in *mdt-15(tm2182)* mutants

Vitamin B12 is the key component of *C. aquatica-induced* developmental acceleration (Watson *et al.* 2014). To determine whether vitamin B12 underlies the *C. aquatica*-induced rescue of the embryonic lethality of *mdt-15(tm2182)* mutants, we grew them on plates supplemented with 50 nM methyl-cobalamin (Me-Cbl), one of the active forms of vitamin B12 (Froese *et al.* 2019). Me-Cbl supplementation strongly rescued the embryonic lethality of *mdt-15(tm2182)* mutants (Fig. 2C). Supplementation with 5 or 500nM of Me-Cbl had a similar effect, and supplementation with 5, 50, or 500 nM adenosyl-cobalamin (Ado-Cbl), another active form of vitamin B12, also effectively rescued embryonic lethality at all concentrations (Fig. 2D). These data show that vitamin B12 levels are essential for normal embryonic viability of *mdt-15* mutants.

### *mdt-15(tm2182)* mutants are sensitive to propionic acid

In *C. elegans*, vitamin B12 is required as a cofactor for two enzymes: the methionine synthase *metr-1;* and the MUT-type isomerase *mmcm-1* (Watson *et al.* 2016). We reasoned that methionine is unlikely to be in short supply in *C. elegans* strains feeding on *E. coli* as food source and thus did not study methionine metabolism. The other *C. elegans* enzyme that utilizes vitamin B12 as a cofactor is *mmcm-1*, which is required to clear propionic acid, a toxic intermediate in the catabolism of odd-chain fatty acids and branched chain amino acids (Watson *et al.* 2016). To determine whether *mdt-15(tm2182)* mutants are sensitive to propionic acid we placed L4 wild-type and *mdt-15* mutant worms grown with or without Me-Cbl on varying concentrations of propionic acid and monitored their survival after 24 hours. Almost 100% of wild-type worms survived in the presence of 20 mM propionic acid, whether or not supplemented with Me-Cbl; in contrast, <2.5% of *mdt-15(tm2182)* mutants survived in the presence of 20 mM propionic acid (Fig. 2E, F). At concentrations between 10 and 20 mM propionic acid, *mdt-15(tm2182)* mutants showed intermediate viability that was significantly rescued by the addition of Me-Cbl (Fig. 2E, F). Thus, loss of *mdt-15* renders *C. elegans* sensitive to propionic acid and this can be rescued by increasing the activity of vitamin B12 dependent propionic acid degradation pathway.

### Unsaturated fatty acids do not rescue the embryonic lethality of *mdt-15* mutants

*mdt-15* is required to express genes for unsaturated fatty acid (FA) synthesis (Taubert *et al.* 2006; Yang *et al.* 2006; Hou *et al.* 2014). To determine whether lack of unsaturated FAs contributes to the embryonic lethality phenotype of *mdt-15(tm2182)* mutants, we grew them on plates containing unsaturated FAs, vitamin B12, or a combination of both. Unlike vitamin B12, unsaturated FAs did not improve embryonic lethality (Fig. 3A). FA supplementation was effective in these experiments, as it rescued the reduced brood size of *fat-6(RNAi)* worms (Fig. 3B), as published (Yang *et al.* 2006).

**Figure 3.**
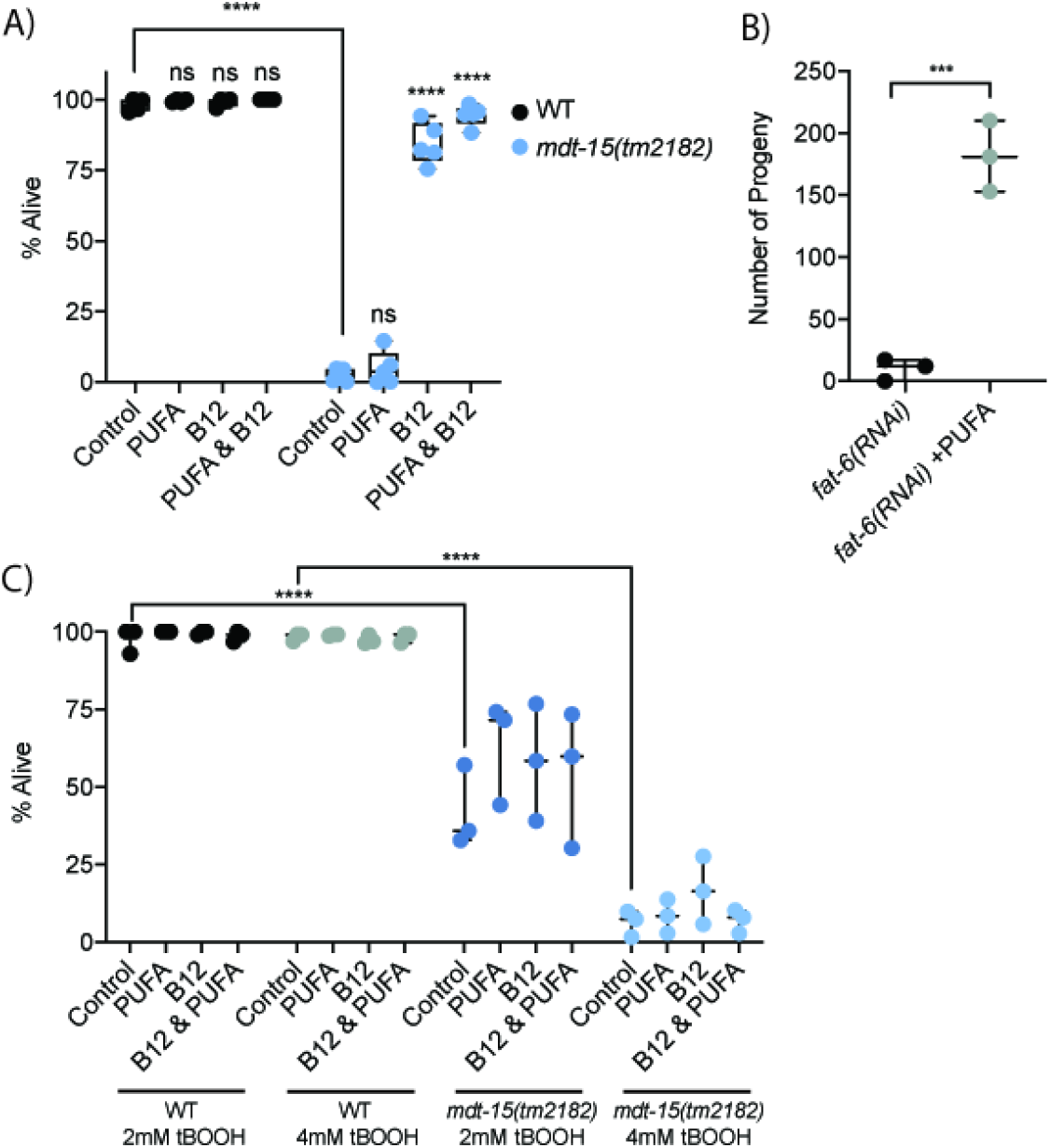
PUFA diets do not rescue embryonic lethality and diets rich in vitamin B12 do not rescue the oxidative stress sensitivity of *mdt-15(tm2182)* mutants. **(A)** The graph shows the average embryonic lethality of wild-type and *mdt-15(tm2182)* mutant worms grown on either *E. coli* OP50, either unsupplemented, or supplemented with methyl-cobalamin, PUFAs, or methyl-cobalamin and PUFAs (n=4-5). ****p<0.001 for indicated comparisons, all other comparisons not significant, as assessed by two-way ANOVA with Tukey correction for multiple comparisons. **(B)** The graph shows the number of progeny laid by *fat-6(RNAi)* worms with and without PUFA supplementations (n=3). ***p<0.001, unpaired Student’s t-test. **(C)** The graph shows the average number of wild-type and *mdt-15(tm2182)* mutant worms that are alive after growth on 2mM or 4mM tBOOH and supplemented with methyl-cobalamin, PUFAs, or methyl-cobalamin and PUFAs, as indicated (n=3). ****p<0.001 for indicated comparisons, ns = not significant, two-way ANOVA with Tukey correction for multiple comparisons.

Besides embryonic lethality, the oxidative stress sensitivity of *mdt-15(tm2182)* mutants is also not rescued by dietary supplementation with PUFAs (Goh *et al.* 2014). We therefore tested whether supplementation with PUFAs, vitamin B12, or a combination of both could rescue the sensitivity of *mdt-15(tm2182)* mutants to the pro-oxidant tert-butyl-hydroperoxide (tBOOH). As before (Goh *et al.* 2014), PUFA supplementation had no effects. Similarly, Me-Cbl supplementation, either alone or in combination with PUFAs, failed to increase tBOOH resistance in *mdt-15(tm2182)* mutant worms (Fig. 3C), suggesting that this phenotype is not related to B12 dependent metabolism.

### *mdt-15* is required to express enzymes in the propionic acid shunt breakdown pathway

Propionic acid is degraded via two pathways in *C. elegans:* the “canonical”, vitamin B12 dependent pathway, and the shunt pathway that is activated and critical when vitamin B12 is low or unavailable (Watson *et al.* 2016). Notably, four of five shunt pathway enzymes (*acdh-1, ech-6, hach-1*, and *hphd-1*), which are repressed when *C. elegans* has sufficient vitamin B12 (MacNeil *et al.* 2013; Watson *et al.* 2014), require *mdt-15* for expression on *E. coli* OP50 (Fig. 1B, Fig. S1B, C). We used RT-qPCR to quantify the expression of these genes in wild-type worms fed *E. coli* OP50, wild-type worms fed *C. aquatica* DA1877, and *mdt-15(tm2182)* mutants fed *E. coli* OP50. This confirmed that *C. aquatica* DA1877 strongly downregulated the expression of *acdh-1, hphd-1, hach-1*, and *ech-6*, as expected (MacNeil *et al.* 2013; Watson *et al.* 2014); in addition, we found that loss of *mdt-15* also downregulated *acdh-1*, *hphd-1*, *hach-1*, and *ech-6* (Fig. 4A). Thus, *mdt-15* is required to activate the expression of some shunt pathway genes in conditions of low vitamin B12 activity.

**Figure 4.**
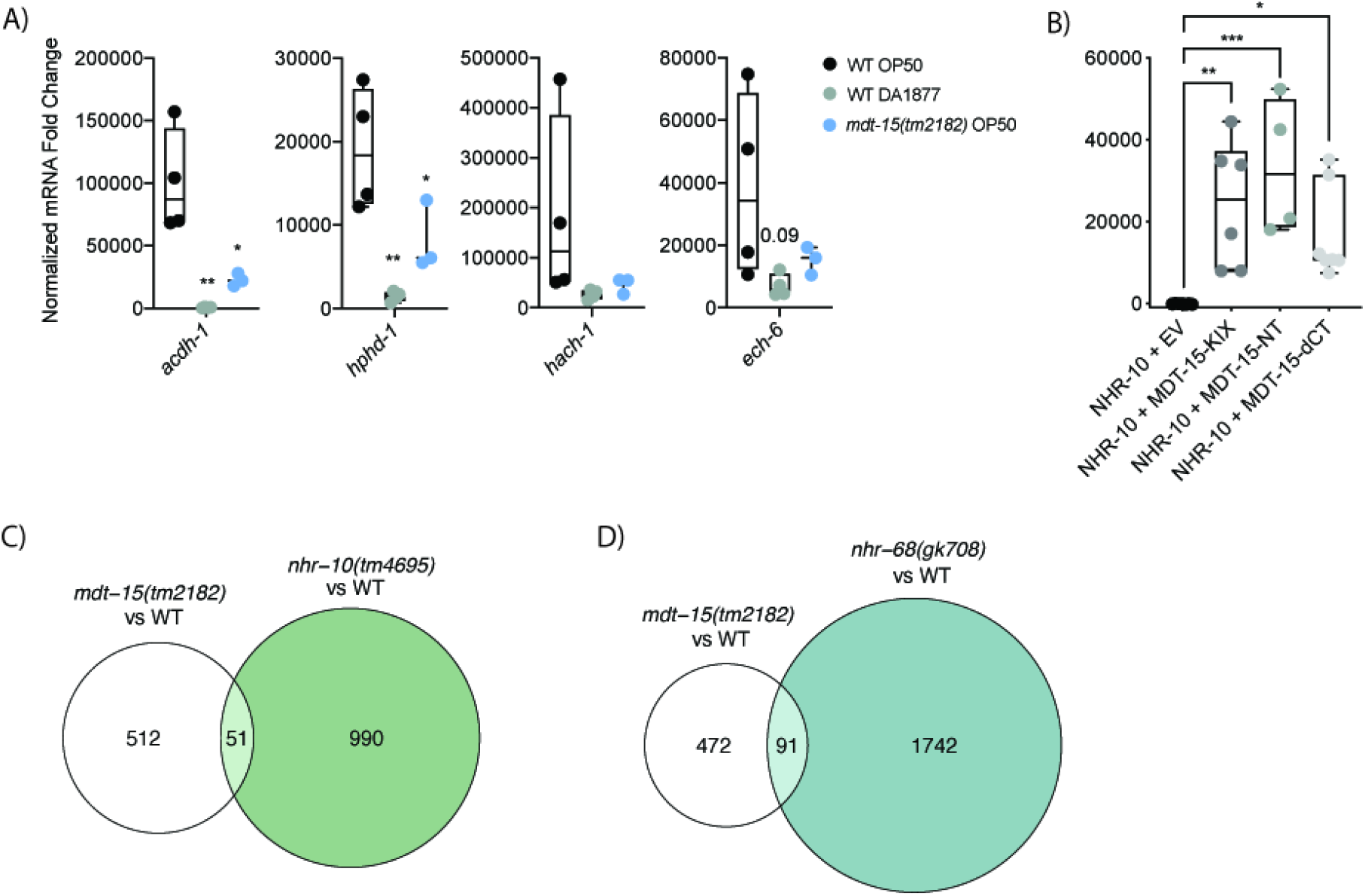
MDT-15 regulates shunt gene expression and binds NHR-10 in Y2H assays. **(A)** The graph indicates relative mRNA levels in L4 wild-type and *mdt-15(tm2182)* mutant worms grown on either *E. coli* OP50 or *C. aquatica* DA1877, as indicated (n=3-4). *p<0.05, **p<0.01, ordinary one-way ANOVA with Tukey correction for multiple comparisons. **(B)** Protein-protein interaction analysis using the Y2H system (n = 4-7). The graph shows the average interaction strength (arbitrary units, A.U.) between an NHR-10 prey and the following baits: empty vector (EV; negative control), MDT-15-KIX (aa-1-124), MDT-15-NT (aa 1-338), or MDT-15ΔCT (aa1-598). Statistical analysis: *p < 0.05, **p < 0.01, ***p < 0.005 vs. NHR-10+EV, Ordinary one-way ANOVA, multiple comparisons, Dunnett correction. **(C)** The Venn diagram shows the overlap of genes downregulated in *mdt-15(tm2182)* and *nhr-10(tm4695)* mutants (Adj.P or PValue<0.05 and logFC<0) compared to WT. Hypergeometric p = 0.00048; platform size = 18605. **(D)** The Venn diagram shows the overlap of genes downregulated in *mdt-15(tm2182)* and *nhr-68(gk708)* mutants (Adj.P or PValue<0.05 and logFC<0) compared to WT. Hypergeometric p = 1.41e-06; platform size = 18605. Lists of genes in each data set are shown in Supplementary Tables S2 and S6-7.

### MDT-15 and NHR-10 co-regulate propionic acid shunt breakdown genes

The nuclear hormone receptors NHR-10 and NHR-68 regulate propionic acid shunt breakdown genes (Bulcha *et al.* 2019). We previously showed that NHR-68 doesn’t bind MDT-15 in yeast-two hybrid (Y2H) assays (Taubert *et al.* 2006). In contrast, we and others detected NHR-10 binding to MDT-15 in Y2H screens (Arda *et al.* 2010; Reece-Hoyes *et al.* 2013). Quantification of binding using Y2H assays revealed that an NHR-10 bait protein indeed strongly bound to several MDT-15 prey proteins (Fig. 4B). Moreover, a construct containing only the KIX-domain of MDT-15 (aa 1-124), previously characterized as an NHR binding domain in MDT-15 (Taubert *et al.* 2006; Goh *et al.* 2014; Shomer *et al.* 2019), was sufficient to mediate this interaction (Fig. 4B). Overlap of gene sets dependent on NHR-10 and NHR-68 (Bulcha *et al.* 2019) and MDT-15 (this study) revealed that the genes controlled by each of these transcriptional regulators overlap substantially (Fig. 4C-D), with the shunt genes amongst the most strongly downregulated genes in all of the mutants of these transcriptional regulators (Fig. S1D, E). We conclude that MDT-15 and NHR-10 likely interact physically and functionally to control the expression of propionic acid shunt breakdown and other genes *in vivo*, acting upstream of NHR-68.

## Discussion

Vitamin B12 is an essential cofactor that is only synthesized by some species of bacteria. In contrast to the *E. coli* OP50 diet normally fed to *C. elegans* in the laboratory, a diet of *C. aquatica* DA1877 contains higher levels of vitamin B12, which influences a number of life history traits in worms, including brood size, developmental rate, and lifespan. Here, we show that vitamin B12 is essential for embryonic viability in *C. elegans* worms carrying a mutation in the *mdt-15* gene. Dissecting this requirement further, we find that *mdt-15* is required for the activation of genes in the propionic acid degradation shunt, which acts in parallel to the canonical vitamin B12-dependent propionic acid degradation pathway. Binding analysis with Y2H assays and comparison of mutant transcriptomes suggest that MDT-15 may interact with NHR-10 to regulate shunt gene expression, thus adapting *C. elegans* metabolism in conditions where propionic acid degradation through the canonical vitamin B12-dependent mechanisms is not possible.

The strong embryonic lethality phenotype of the *mdt-15(tm2182)* mutant and its virtually complete rescue by diets rich in vitamin B12 is interesting. Many of the phenotypes of this mutant (larval arrest, fecundity, life span, mobility) are largely rescued by supplementation of worms with unsaturated fatty acids, yet these have no effect on the embryonic lethality phenotype. In line with the phenotype we observed, RNAi and/or mutation of *acdh-1, hach-1, hphd-1*, and *alh-8* also causes embryonic arrest, which for *acdh-1, hphd-1*, and *alh-8* manifests specifically when vitamin B12 is low in abundance (Gönczy *et al.* 2000; Simmer *et al.* 2003; Sönnichsen *et al.* 2005; Watson *et al.* 2016); similarly, *ech-6* RNAi caused larval arrest, albeit not embryonic arrest (possibly due to RNAi efficiency variability). It is not fully clear why vitamin B12, and by inference propionic acid detoxification, is crucial during embryonic development. Possibly, the rapid growth associated with embryonic development leads to transient increases in propionic acid breakdown intermediates, perhaps because specific amino acids or fatty acids are in demand.

We did not test whether the other vitamin B12 activated pathway, methionine synthesis, are affected by *mdt-15* mutation. We reasoned that even if methionine synthesis were low, these worms should receive an adequate methionine supply from their *E. coli* diet. However, methionine synthesis is linked to the one-carbon cycle, in particular the metabolism of folate to its biologically active form, tetrahydrofolate. Interestingly, *mdt-15* mutants express lower levels of the folate transporter *folt-2*, as determined by microarray and RT-qPCR analysis (Taubert *et al.* 2008). However, no overt phenotypes have been reported for *folt-2* depletion by RNAi (Gönczy *et al.* 2000; Simmer *et al.* 2003; Sönnichsen *et al.* 2005), perhaps because *folt-1* and *folt-3* compensate for *folt-2* loss.

Our study adds to the growing list of specific gene regulatory programs that are implemented by partnerships between MDT-15 and NHRs (and other TFs). For example, we showed that NHR-49 and MDT-15 interact physically and regulate the expression of numerous fatty acid metabolism genes, which in turn affects numerous phenotypes of *mdt-15* mutants, including viability, life span, fecundity, mobility, and others (Taubert *et al.* 2006; Yang *et al.* 2006). Recent studies suggest that NHR-49 and MDT-15 also interact to promote resistance to several stresses, including starvation, oxidative stress, hypoxia, and pathogen resistance (Goh *et al.* 2018; Hu *et al.* 2018; Dasgupta *et al.* 2020; Hummell *et al.* 2021; Wani *et al.* 2021; Doering *et al.* 2022). MDT-15 also binds NHR-86 (Reece-Hoyes *et al.* 2013), and both factors regulate the expression of innate immune response genes and ensure survival in response to infection with the pathogen *Pseudomonas aeruginosa* (Peterson *et al.* 2019). Furthermore, an interaction between MDT-15 and HIZR-1 (aka NHR-33) promotes the expression of heavy metal response genes through the High-zinc activated regulatory element (Roh *et al.* 2015; Shomer *et al.* 2019). The data presented here suggest that NHR-10 and MDT-15 interact to transcriptionally induce propionic acid degradation shunt genes when propionic acid levels are high and/or vitamin B12 levels are low. Interestingly, HIZR-1 directly binds the essential metal zinc as well as non-essential, toxic cadmium (Warnhoff *et al.* 2017; Earley *et al.* 2021), and HIZR-1 binding to MDT-15 is strongly increased in response to such ligand binding (Shomer *et al.* 2019). This suggests that metabolites linked to the propionic acid degradation pathway may also be ligands for NHRs such as NHR-10, although binding of an organic acid with a carbon length as short as three to an NHR has not yet been shown to our knowledge. It is possible that upstream metabolites, such as branched-chain amino acids and longer odd-chain fatty acids, act as ligands for NHR-10.

In summary, we report a novel interaction between the transcriptional regulator MDT-15 and vitamin B12, an essential micronutrient. We hypothesize that, when vitamin B12 levels are low, and/or propionic acid levels are high, NHR-10 and MDT-15 interaction is stimulated, which in turn allows increased expression of the shunt genes and permits normal embryonic development despite low vitamin B12 availability. It will be interesting to determine whether this mechanistic role for MDT-15 and HNF4-like nuclear receptors is conserved in mammals, especially in view of shunt gene activation by exogenous propionic acid in culture liver cancer cell lines (Watson *et al.* 2016).

## Data availability

The data underlying this article are available in the article, in its online supplementary material, and at Gene Expression Omnibus online (GSE220955). All reagents are available upon request. Supplemental material is available at G3 online.

## Acknowledgements

We thank Taubert lab members and Amy K. Walker (U Mass Medical School, Worcester, MA, USA) for comments on the manuscript.

## Figure legends

**Supplementary Figure S1. Transcriptomes from specific gene inactivations. (A)** The scatter plot shows the correlation of differentially expressed genes (Adj.P or PValue<0.05) in *mdt-15(tm2182)* mutant worms and *mdt-15(RNAi)* worms. X and Y axes represent logFC. Pearson r=0.66; p=0. **(B-E)** The volcano plots show the expression of all detected genes in **(B)** *mdt-15(RNAi) vs.* EV, **(C)** *mdt-15(tm2182) vs.* WT, **(D)** *nhr-10(tm4695) vs.* WT, and **(E)** *nhr-68(gk708) vs.* WT worms. The shunt genes and *mdt-15* are highlighted. X-axis, logFC; Y-axis, -log_10_(PValue or Adj.P, as indicated). Black, PValue or Adj.P <0.05; grey, PValue or Adj.P ≥0.05; blue, highlighted and significantly downregulated; purple, highlighted but not significant.

## Supplementary Tables

**Supplementary Table S1.** The table shows the number of unique significantly regulated DEGs from each transcriptome dataset. Significance is defined as PValue or Adj.P <0.05. Up- and down-regulation are defined as logFC>0 and logFC<0, respectively. Study type, relevant GEO accession numbers, and source publications are also listed.

**Supplementary Table S2.** List of unique significantly regulated DEGs (Adj.P<0.05) in *mdt-15(tm2182)* mutants *vs.* WT worms.

**Supplementary Table S3.** List of unique significantly regulated DEGs (PValue<0.05) in WT worms fed *mdt-15(RNAi) vs.*EV RNAi control.

**Supplementary Table S4.** List of unique significantly regulated DEGs (PValue<0.05) in WT worms fed *C. aquatica* DA1877 *vs.*HT115.

**Supplementary Table S5.** List of unique significantly regulated DEGs (PValue<0.05) in WT worms fed OP50 supplemented with B12 *vs.* OP50 control.

**Supplementary Table S6.** List of unique significantly regulated DEGs (PValue<0.05) in *nhr-10(tm4695)* mutants *vs.* WT worms.

**Supplementary Table S7.** List of unique significantly regulated DEGs (PValue<0.05) in *nhr-68(gk708)* mutants *vs.* WT worms.

